# Stronger brain responses to acute stress reflect greater everyday stress variability

**DOI:** 10.64898/2026.08.26.747278

**Authors:** Madeleine Kördel, Anne Kühnel, Ann-Christin S. Kimmig, Sandra Beinbauer, Lydia Kogler, Inger Sundström-Poromaa, Melanie Henes, Nils B. Kroemer

**Author notes:** **Corresponding author:** Prof. Dr. Nils B. Kroemer, Venusberg Campus 1, 53127 Bonn, Germany.

## Abstract

Laboratory stress tasks are widely used to assess individual differences in acute stress reactivity, yet it remains unclear how these responses correspond to stress experienced in everyday life. Here, we combined the Montreal imaging stress task (MIST) with ecological momentary assessment (EMA) over three months to assess acute and everyday stress in 67 healthy women. Greater within-person variability in everyday stress, but not average stress levels, were associated with stronger overall stress-related brain responses (*b* = 0.73, *p* = .039), with a whole-brain association particularly evident in the bilateral caudate (*r_ROI_* = .32, *p_cluster.FWE_* < .001). Greater everyday stress variability was also associated with stronger stress-related functional connectivity between the ventromedial prefrontal cortex (vmPFC) and parietal and posterior medial regions (*p_cluster.FWE_* < .001). We conclude that acute neural stress responses relate more closely to fluctuations in perceived stress than to how stressed an individual feels on average. This suggests that laboratory stress tasks capture acute stress responsivity that is distinct from average stress exposure, highlighting the importance of considering what these tasks measure when interpreting individual differences in acute stress responses.

## Introduction

Stress is experienced often in everyday life and plays an important role in mental and physical health. Acute stress responses are generally adaptive as they mobilize cognitive, autonomic, and endocrine resources to meet immediate environmental demands (Chrousos, 1992; Habib et al., 2001; Sapolsky et al., 2000). Their coordination involves limbic and prefrontal brain regions, including the hippocampus and prefrontal cortex, which contribute to the modulation of the neuroendocrine stress response (Dedovic et al., 2005; Herman et al., 2005; Pruessner et al., 2008). Among these regions, the ventromedial prefrontal cortex (vmPFC) integrates cognitive, emotional, and physiological information to support adaptive stress responses (Ginty et al., 2019; Roy et al., 2012), while the hippocampus contributes to contextual processing and the negative feedback regulation of the hypothalamic-pituitary-adrenal (HPA) axis (Elbau et al., 2018; Pruessner et al., 2008). These acute stress responses are typically examined using standardized laboratory stress paradigms. However, it remains unclear whether individual differences observed under laboratory conditions correspond to how individuals experience and respond to stress in their everyday lives. Addressing this gap is relevant for understanding individual differences in vulnerability to stress-related mental health problems.

Acute laboratory stress can lead to changes in stress-related brain responses across several brain regions, including those of the salience and executive networks (Berretz et al., 2021; Henze et al., 2020; Kuhn et al., 2023; Kühnel et al., 2022). Functional connectivity studies further indicate that acute stress alters vmPFC coupling with regions involved in salience, cognitive control and stress regulation (Ginty et al., 2019; Vaisvaser et al., 2016). Extending these findings, recent research combining laboratory stress induction with ecological momentary assessment (EMA) shows that greater salience network activation during laboratory stress is associated with stronger affective responses to everyday stressors (Tutunji et al., 2025). Complementary longitudinal work adds to this by showing that neural responses to acute psychosocial stress in the amygdala and hippocampus are associated with increases in perceived daily life stress over the following year (Giglberger et al., 2023). Together, these findings suggest that acute stress-related brain responses may reflect individual differences in real-life stress sensitivity and longer-term stress vulnerability.

Repeated or ongoing stress in everyday life may also be associated with neural responses to laboratory stress. In one of the few studies directly addressing this relationship, higher daily stress has been linked to reduced hippocampal activation and stronger connectivity between the hippocampus and the vmPFC during acute stress (Ren et al., 2022). Neuroimaging studies further suggest that high chronic stress, compared to low, is associated with less pronounced changes in vmPFC-amygdala and vmPFC-hippocampus connectivity from acute stress to recovery (Chen et al., 2026). Related evidence from clinical populations shows that individuals with major depressive disorder exhibit reduced vmPFC activation (Ming et al., 2017) and reduced prefrontal-amygdala connectivity (Hossein et al., 2023) to acute stress compared with healthy control participants. Thus, the available research indicates that acute stress-related brain responses may vary with higher daily stress and in clinical populations, although results linking stress experienced repeatedly in everyday life to laboratory stress in healthy individuals remain scarce.

So far, existing studies have typically assessed everyday stress using relatively sparse measurements collected over short periods or at isolated time points, or have relied on retrospective questionnaires administered during laboratory sessions (De Calheiros Velozo et al., 2023; Giglberger et al., 2023; Ren et al., 2022; Tutunji et al., 2025). Additionally, in human neuroimaging research, the endocrine status of women, including oral contraceptive (OC) use, is rarely considered (Taylor et al., 2021), despite evidence that endogenous and exogenous sex hormones can modulate affective stress reactivity (Bürger et al., 2025), HPA axis functioning (Barel et al., 2018; Kördel et al., 2025) and stress-related brain responses (Albert et al., 2015; Sharma et al., 2020). As a result, little is known about how brain responses to acute stress relate to stress dynamics experienced repeatedly in everyday life over extended periods or how this relationship may vary with factors that contribute to individual differences in stress responses in women.

To address these gaps, the present study examines how everyday stress, assessed with EMA and tracked over three months, relates to brain responses to an acute psychosocial stressor in a sample of healthy young women. We combine a standardized functional magnetic resonance imaging (fMRI) stress paradigm with multimodal assessments of subjective stress and autonomic responses, and we examine both regional BOLD responses and stress-induced changes in functional connectivity, with a focus on vmPFC-centered circuits. The integration of fMRI with EMA allows us to bridge laboratory-based stress reactivity and everyday stress experiences. We expect acute stress responses to be associated with self-rated stress experiences in everyday life and observed that greater variability, but not average levels, in everyday stress were linked to stronger stress-related brain responses. Our findings suggest that acute laboratory stress paradigms capture acute stress responsivity or sensitivity that is separable from average stress exposure. Distinguishing between these aspects of stress is important for understanding what laboratory stress tasks measure and for interpreting individual differences in acute stress responses.

## Methods

The data was part of a larger, ongoing study specifically examining the association between OC and stress reactivity registered at clinicalTrials.gov (NCT06223126). Here, we focus on the association between acute and everyday stress independent of the intervention effects, which will be reported after completion of the study.

### Ethics Statement

The study was approved by the Ethics Committee of the Faculty of Medicine at the University of Tübingen (412/2023BO1). All procedures were carried out in accordance with the Declaration of Helsinki. Written informed consent was obtained from all participants before the experiment.

### Participants

In the present analysis, we included data collected until the data freeze in February 2026. The final sample comprised 67 healthy women (*M_age_*: 25.1 ± 4.3 years, range: 19-38, *M_BMI_*: 22.7 ± 2.7 kg/m^2^, range: 18.4-33). Of these, 53 used combined OC and 14 were assessed during the early follicular phase (days 1-7). A total of 75 participants were initially enrolled, of whom 6 withdrew after the screening session and two were excluded due to missing fMRI data. After the first fMRI session, OC use status changed in a subset of participants during the subsequent EMA period. The 14 naturally cycling women started OC use (OC starters), 19 OC users stopped OC use (OC stoppers), and 34 participants continued OC use (OC long-term). Exclusion criteria included mental, neurological, or significant medical conditions, pregnancy or breastfeeding within the past year, regular medication use (except OC) and MRI incompatibility. They received monetary compensation of 100€ upon completion of the study.

### Experimental procedure

Participants completed three laboratory sessions over three months (Figure 1). The present study included data from the screening session (S0), the first fMRI session (S1), and the subsequent EMA period. In S0, participants arrived after an overnight fast (>12 h) between 7 a.m. and 11 a.m. and were screened for mental disorders. Sociodemographic information and MRI compatibility were assessed using standardized questionnaires as part of the screening procedure to identify exclusion criteria. A fasting blood sample was collected to determine baseline levels of cortisol. After that, we invited participants for two fMRI sessions (S1 and S2), the first before changing OC use and the second approximately 3 months later. Both fMRI sessions were conducted in the afternoon or evening on weekdays (*M_start time_* = 3:54 p.m. ± 86.4 min). At each fMRI session, participants underwent a high-resolution structural scan and a resting-state scan, followed by the Montreal Imaging Stress Task (MIST; adapted from Dedovic et al., 2005), a standardized psychosocial stress paradigm. Heart rate was continuously recorded during the resting-state scan and MIST. Participants provided subjective stress ratings at six time points, including baseline, before and after the stress task inside the scanner, and approximately 40, 60 and 90 minutes after stress onset outside the scanner. Current stress was rated on a 10-point visual analogue scale (VAS) anchored at 1 (“not at all”) and 10 (“very much”). Additional biological samples (i.e., blood, saliva, and hair) and questionnaire data were collected but are not reported here.

**Figure 1.**
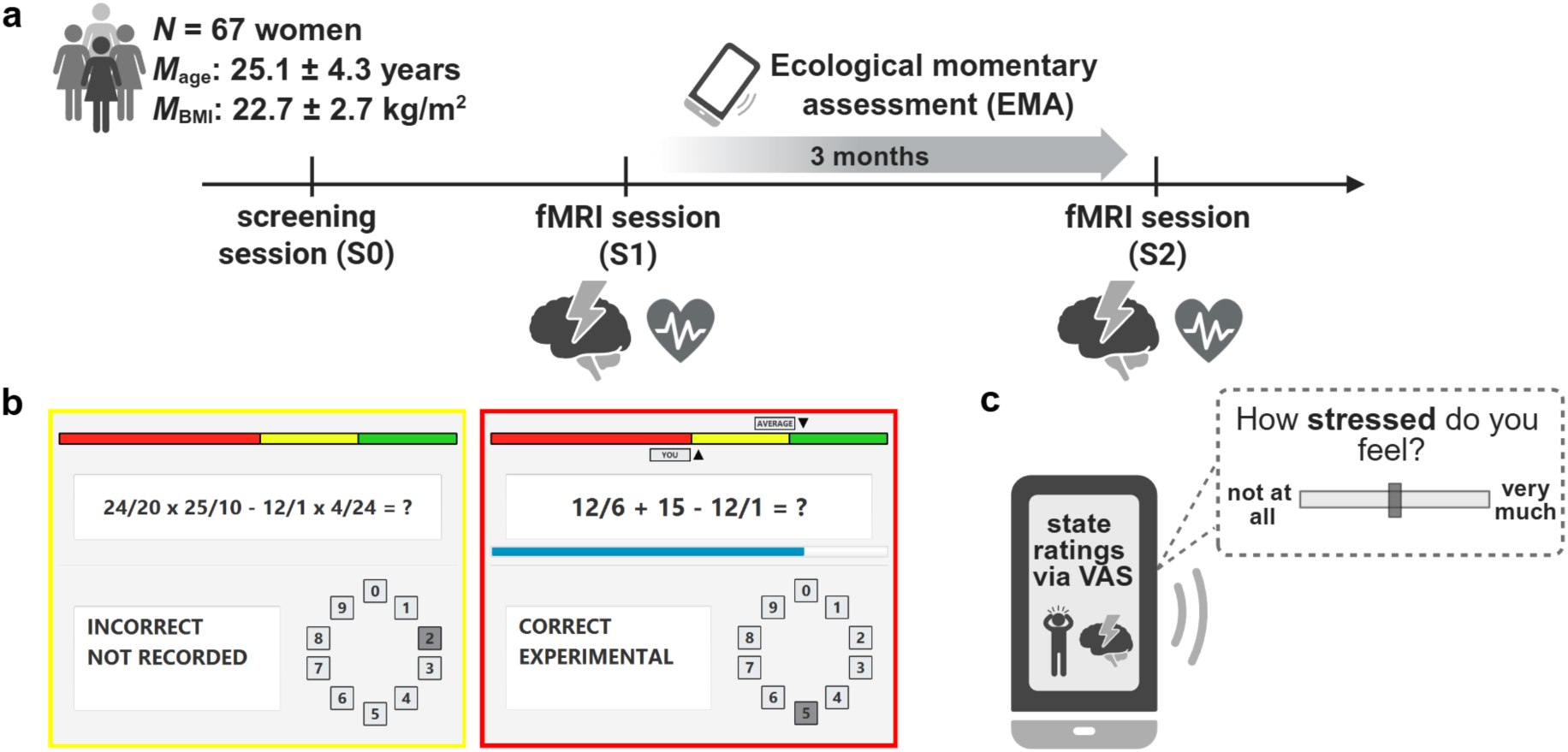
Summary of the study. **a:** Timeline of the experimental procedure showing three laboratory visits (S0-S2). Participants completed two fMRI sessions (S1 and S2) and ecological momentary assessments (EMA) in the three-month interval between them. Heart rate was continuously recorded during S1 and S2, and subjective stress was assessed at six time points throughout these two sessions. **b:** The Montreal Imaging Stress Task (MIST; adapted from Dedovic et al., 2005) was administered during both fMRI sessions, with the control condition depicted in yellow and the stress condition in red. **c:** Participants rated their momentary stress once per day using a visual analogue scale (VAS) ranging from “not at all” to “very much.” Figure created with BioRender.com.

Following the first MRI session (S1), participants completed EMA over three months. Assessments were administered two to three times per week on randomly selected days to capture current stress ratings in everyday life (*M* = 26.5 ± 4.78 responses per participant; range: 9–35; mean compliance rate of 88.4% ± 15.9%). Participants rated their perceived stress “at the moment” on a VAS ranging from 1 (“not at all”) to 10 (“very much”), reflecting momentary stress. Affective states were assessed with reference to the respective day using several items, including “How depressed did you feel today?” and “How satisfied did you feel today?”, with responses provided on a VAS ranging from 1 (“not at all”) to 100 (“very much”). EMA prompts were delivered to participants’ smartphones at 6:00 p.m. and responses could be submitted until midnight.

### Stress task

The MIST is a frequently used psychosocial stress paradigm in neuroimaging settings (Dedovic et al., 2005), in which arithmetic tasks are presented to generate performance demands. These tasks are solved either without time pressure and without social evaluation (control condition) or with time pressure and social evaluation (stress condition), while the initially selected difficulty level remained the same for both conditions. The modified version of the MIST used in this study consisted of a block design with three imaging runs, encompassing Control (∼61s) and Stress (∼90s) conditions. Each condition is presented twice per run (order: control-stress-control-stress), resulting in a total time of 5 minutes per run. Visual feedback during the task is presented as ’TIME OUT’, ’FALSCH’ (wrong), and ’RICHTIG’ (correct). During the stress condition, a time bar and a performance bar appear, and individual performance is compared to a fictional control group. To increase stress, the study instructor provides scripted negative feedback between each run. Participants responded to the task via a button box, and the task was practiced outside the scanner in a training session to familiarize the participants with the procedure.

### Heart rate

During a resting-state scan and the MIST, we recorded pulse waveforms using an MRI-compatible pulse oximeter (version 5.0.8; BIOPAC Systems, Inc., 2024) attached to the left index finger. Heart rate (HR) and heart rate variability (HRV) were analyzed using Kubios HRV software (version 4.1.2.1; Kubios Oy, 2023). Analyses followed the standard configuration of the software, including artifact-corrected interpolation and time- and frequency-domain metrics. R-peak detection was visually inspected, and artifacts were manually removed when necessary. We excluded data segments with more than 5 % artifacts. HR data from 12 participants were excluded from the analysis because more than 50% of heart rate measurements across the MIST control and stress blocks were missing.

### fMRI data acquisition and preprocessing

fMRI data were acquired on a 3T Magnetom Prisma scanner (Siemens Medical Solutions, Erlangen, Germany) using a 64-channel head coil. Functional images were collected with a multiband gradient-echo echo-planar imaging (EPI) sequence (time of repetition [TR] = 1400 ms, time of echo [TE] = 30 ms, flip angle = 65°; matrix = 110 × 110; field of view = 220 × 220 mm²). For each MIST run, 216 volumes were acquired with whole-brain coverage (68 slices; slice thickness = 2 mm; voxel size = 2.0 × 2.0 × 2.0 mm³; no inter-slice gap). Functional data were preprocessed using fMRIPrep (version 23.2.0) (Esteban et al., 2019). Preprocessing included slice-timing correction, head motion correction using rigid-body realignment, and susceptibility distortion correction using a fieldmap-based approach with a phase-difference map. Functional images were co-registered to the individual T1-weighted anatomical image using boundary-based registration with six degrees of freedom and spatially normalized to standard space (full description, see SI). Head-motion parameters and physiological noise regressors derived from white matter and cerebrospinal fluid signals (CSF) were estimated for subsequent denoising in first-level analyses.

## fMRI data analysis

### First-level functional activation analysis

Functional MRI data were analyzed using Statistical Parametric Mapping (SPM12; https://www.fil.ion.ucl.ac.uk/spm/software/spm12/). For the first-level analysis, we estimated task-related activity across the three MIST runs using a general linear model. We included a regressor for the stress blocks (duration 90s) and one regressor capturing button presses (modeled as events with zero duration), both convolved with the canonical hemodynamic response function (with the control condition as implicit baseline). To account for the longer task blocks, we used a high-pass filter of 256 seconds. Motion parameters and average white matter and CSF signals were included as nuisance regressors.

### First-level functional connectivity (gPPI) analysis

To examine stress-induced changes in connectivity, we used generalized psychophysiological interaction (gPPI) analyses implemented in the gPPI toolbox in SPM12 (McLaren et al., 2012). As a seed region, we chose the vmPFC because of its central role in adaptive stress responses and in modulating stress-related network activity (Ginty et al., 2019; Ren et al., 2022; Roy et al., 2012). The vmPFC seed mask was derived from the Neurosynth resting-state functional connectivity map centered at MNI coordinates x = 0, y = 36, z = -14 and binarized in SPM12 using ImCalc with a threshold of > 0.63 (https://www.neurosynth.org/locations/?x=0&y=36&z=-14). We then estimated gPPI models that included the interaction between the vmPFC time course and the stress condition to quantify changes in vmPFC connectivity by comparing the stress condition with the control condition.

### Second-level analysis

The contrast images (stress vs. control) from the first-level models (activation and vmPFC connectivity) were then used for the second-level whole-brain analyses using multiple regression models. In these models, the average self-reported stress (mean) and its variability (SD) across the 3-month EMA phase were regressors of interest, with age, BMI, and OC use group (OC use vs. non-use) in the first MRI session included as nuisance covariates.

To obtain individual summary measures of stress-related activation and deactivation in task-positive and task-negative networks, we first derived consensus masks from the first-level stress vs. control contrast images using a leave-one-subject-out procedure. For each participant, a voxelwise one-sample *t*-test against zero was performed on the contrast images of all remaining participants using a threshold of *p* < .001. Voxels that showed significant positive or negative effects consistently across all leave-one-out iterations were retained in the final activation and deactivation consensus masks, respectively. For each participant, contrast estimates were then extracted from both masks and averaged across voxels to obtain mean activation and mean deactivation values for subsequent analyses. As a post hoc analysis, we examined whether associations with EMA stress variability were specific to individual regions or reflected a more general modulation of the task-positive and task-negative response. To this end, we examined the associations of mean and SD of EMA stress with individual mean activation and deactivation values using separate linear regression models. For visualization of the whole-brain findings, contrast estimates were additionally extracted from regions showing significant effects using the Harvard-Oxford extended atlas (Desikan et al., 2006; Pauli et al., 2018; Teckentrup et al., 2021) and averaged across hemispheres where applicable.

To examine whether effects were predominant in specific functional networks, we assigned voxelwise *t*-values to the seven canonical Yeo functional networks (Yeo et al., 2011). Subcortical voxels not covered by the cortical parcellation were parcellated using the Harvard-Oxford extended atlas (Desikan et al., 2006; Pauli et al., 2018; Teckentrup et al., 2021) and classified as either diencephalon, brainstem, or midbrain (Müller et al., 2022). Mean stress vs. control contrast estimates were extracted from each participant’s first-level contrast image within each network and subcortical category, following a previously described approach (Kühnel et al., 2020). These estimates were tested against zero using two-sided one-sample *t*-tests. Associations with EMA stress variability were examined using separate linear regression models for each network and subcortical category, including mean and SD of EMA stress as predictors of interest and grand mean-centered age, BMI, and OC use as covariates. The same procedure was applied to the gPPI analyses using the individual stress vs. control connectivity contrast images. *p*-values were corrected across the ten categories using Benjamini-Hochberg false discovery rate (FDR) correction. Whole-brain results were corrected for multiple comparisons using cluster-level family-wise error (FWE) correction at *p* < .05, with an initial voxel-level threshold of *p* < .001. The corresponding MATLAB/R code and data are available on GitHub (https://github.com/neuromadlab/Stress_EMA_fMRI). Brain contrast images for the second-level whole-brain and functional connectivity results are uploaded on NeuroVault.org (https://identifiers.org/neurovault.collection:24307).

### Statistical analysis and software

As manipulation checks, we first tested whether the stress induction was successful. We examined HR changes in response to the stress task using a linear mixed-effects model (LME), with HR as the dependent variable and condition (resting-state, control, and stress) as a fixed effect (165 observations from 55 participants due to missing data). Changes in self-reported stress over the course of the task were examined in a second LME, with stress ratings as the dependent variable and measurement timepoint (baseline, stress onset, and 25, 45, 60, and 90 minutes after stress onset) as a fixed effect (402 observations).

For the fMRI session, mean HR was calculated separately for the resting-state scan, across the entire MIST (mean across all three runs), and across the stress and control conditions of the MIST. Resting-state HR served as the baseline measure. The task-induced HR response was quantified as the difference between mean HR across the entire MIST and mean HR during the resting-state scan. The task-induced subjective stress response was calculated by subtracting the pre-stress rating immediately before stress onset from the post-stress rating immediately after the stress task. For EMA stress, participant-level mean and standard deviation (SD) scores were calculated across ratings of the “momentary stress” item to characterize average stress levels and stress variability, respectively.

Our main analyses examined whether individual differences in the subjective and autonomic stress response to the laboratory stress task were associated with EMA stress. We used separate LMEs with EMA stress ratings as the outcome and either the task-induced subjective stress response or the task-induced HR response as predictors. In both models, grand mean-centered EMA mood state was additionally included as a predictor to test whether the respective variable of interest explained stress beyond the participant’s mood state. Mood state was computed as a composite of satisfied and depressed ratings by subtracting the negative mood item (depressed) from the positive mood item (satisfied) to capture mood along a positive-to-negative valence dimension, consistent with previous EMA analyses (Kaduk et al., 2026; Kördel et al., 2025).

As sensitivity analyses, we tested whether OC group was associated with individual differences in average EMA stress levels and EMA stress variability. We used two separate linear regression models, with the mean and SD of EMA stress as the respective dependent variables and OC group as the predictor.

Analyses included data from the screening session, first fMRI session, and EMA phase, while data from S2 was not included. All models used grand mean-centered age and BMI as well as effect-coded OC use at session 1 (0.5 = use, -0.5 = no use; except for the OC sensitivity analyses) as covariates. LME models included a random intercept for participant and random slopes for within-person predictors where applicable. Significance of model effects was evaluated using ANOVAs, and significant effects with more than two levels were followed up using Tukey-adjusted estimated marginal means. We conducted all statistical analyses in RStudio (2026.7.1.147; Posit team, 2026). LME models were estimated using lmerTest (Kuznetsova et al., 2017) with the Satterthwaite correction for degrees of freedom and post-hoc comparisons were performed using the emmeans package (Lenth & Piaskowski, 2025). We considered α ≤ .05 as significant. Data were visualized using ggplot2 (Wickham, 2016), ggsignif (Constantin & Patil, 2021) and ggdist (Kay, 2024, 2025). For brain images, we used Mango image processing software (v4.1; Research Imaging Institute, UTHSCSA).

### Results Stress-task induces multimodal stress response

We first assessed whether the stress task elicits the expected increases in self-reported stress, HR and brain response. On a subjective and physiological level, the task successfully induced acute stress. HR differed significantly across conditions (Figure 2a; *F*(2, 108) = 91.78, *p* < .001). Post hoc comparisons showed that mean HR significantly increased from the resting-state scan to the control (*b* = 17.47, *t*(108) = 11.29, *p* < .001) and the stress condition (*b* = 18.77, *t*(108) = 12.13, *p* < .001). However, HR did not differ significantly between the control and stress conditions (*b* = 1.29, *t*(108) = 0.84, *p* = .681). Likewise, subjective stress ratings differed significantly across the six assessed timepoints (Figure 2b; *F*(5, 330) = 121.86, *p* < .001), with ratings increasing significantly in response to the stress task. Higher ratings were reported immediately after the task (approximately 25 min after stress onset) compared to immediately before (post hoc test: *b* = 4.54, *t*(330) = 17.05, *p* < .001). Ratings then declined across the recovery period, dropping significantly at each subsequent timepoint relative to the post-task peak (all *p* < .001). Ratings remained significantly elevated above pre-task baseline through 45 min post-task (*b* = 0.94, *t*(330) = 3.53, *p* = .006), before falling below pre-task baseline levels by 60 and 90 minutes after the task (both *p* < .020).

**Figure 2.**
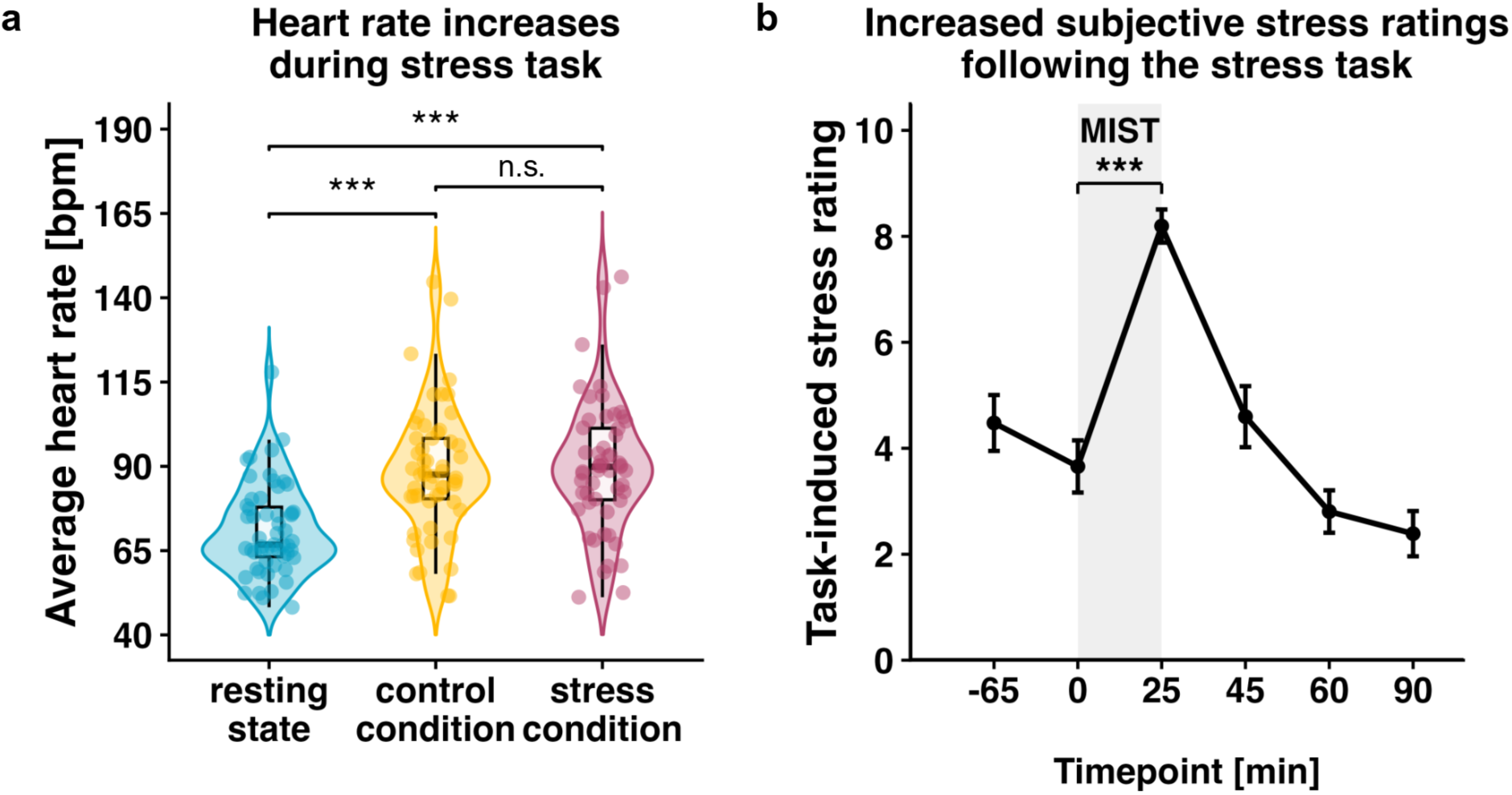
The MIST induces subjective and autonomous stress responses. **a**: Violin plots show average heart rate (bpm; Boxplots indicate the interquartile range and median; individual points depict participants with ≥50% complete heart-rate data, *N* = 55), across the resting-state scan and the control and stress conditions of the MIST. The mean heart rate increased significantly during the control and stress conditions (\*\*\**p* < .001, n.s. = not significant). **b:** Line plot of mean task-related stress ratings (± SEM) across timepoints relative to the MIST (shaded area). Ratings peaked immediately after the stress task (25 min) and returned toward baseline during recovery. The MIST significantly increased subjective stress compared to pre-task ratings (\*\*\**p* < .001). These findings indicate effective stress induction at subjective and physiological levels.

Given the role of cortisol in stress regulation, we examined whether fasting cortisol levels obtained during the screening session (S0) were associated with autonomic and subjective responses measured during the fMRI session and EMA. Higher cortisol levels were associated with higher mean HR across the stress task (Figure S1; *r* = .28, *p* = .027). Cortisol was also not associated with resting-state HR, task-induced subjective stress, mean EMA stress, or variability in EMA stress (all *p* ≥ .279).

At the whole-brain level, the stress condition elicited greater activity than the control condition in the anterior cingulate cortex (*t_max_* = 8.54), anterior insula (*t_max_* = 7.63), thalamus (*t_max_* = 6.54), middle frontal gyrus (*t_max_* = 9.37), middle occipital gyrus (*t_max_* = 11.30) and middle temporal gyrus (*t_max_* = 14.57, all *p_cluster.FWE_* < .001). As expected, activation in the default mode network, including the angular gyrus (*t_max_* = 11.23), precuneus (*t_max_* = 11.13) and inferior parietal gyrus (*t_max_* = 9.12) as well as the posterior insula (*t_max_* = 11.77), posterior orbitofrontal cortex (OFC; *t_max_* = 9.10), bilateral nucleus accumbens (NAc; *t_max_* = 8.21), amygdala (*t_max_* = 6.33) and parahippocampal gyrus (*t_max_* = 8.08, all *p_cluster.FWE_* < .001; Fig. 3a-b, Table S1), was lower during stress vs. control. At the network level, this was reflected by increased activity in the dorsal attention network (*t*(66) = 4.49, *p_FDR_* < .001) and midbrain (*t*(66) = 2.38, *p_FDR_* = .040) (Yeo et al., 2011) and decreased activity in the default mode (*t*(66) = -3.47, *p_FDR_* = .003), limbic (*t*(66) = -6.38, *p_FDR_* < .001) and somato-motor networks (*t*(66) = -2.61, *p_FDR_* = .028) during stress (Figure 3c). No significant effects were observed in the visual, ventral attention, frontoparietal, diencephalon, or brainstem categories after FDR correction (all *p_FDR_* ≥ .074).

**Figure 3.**
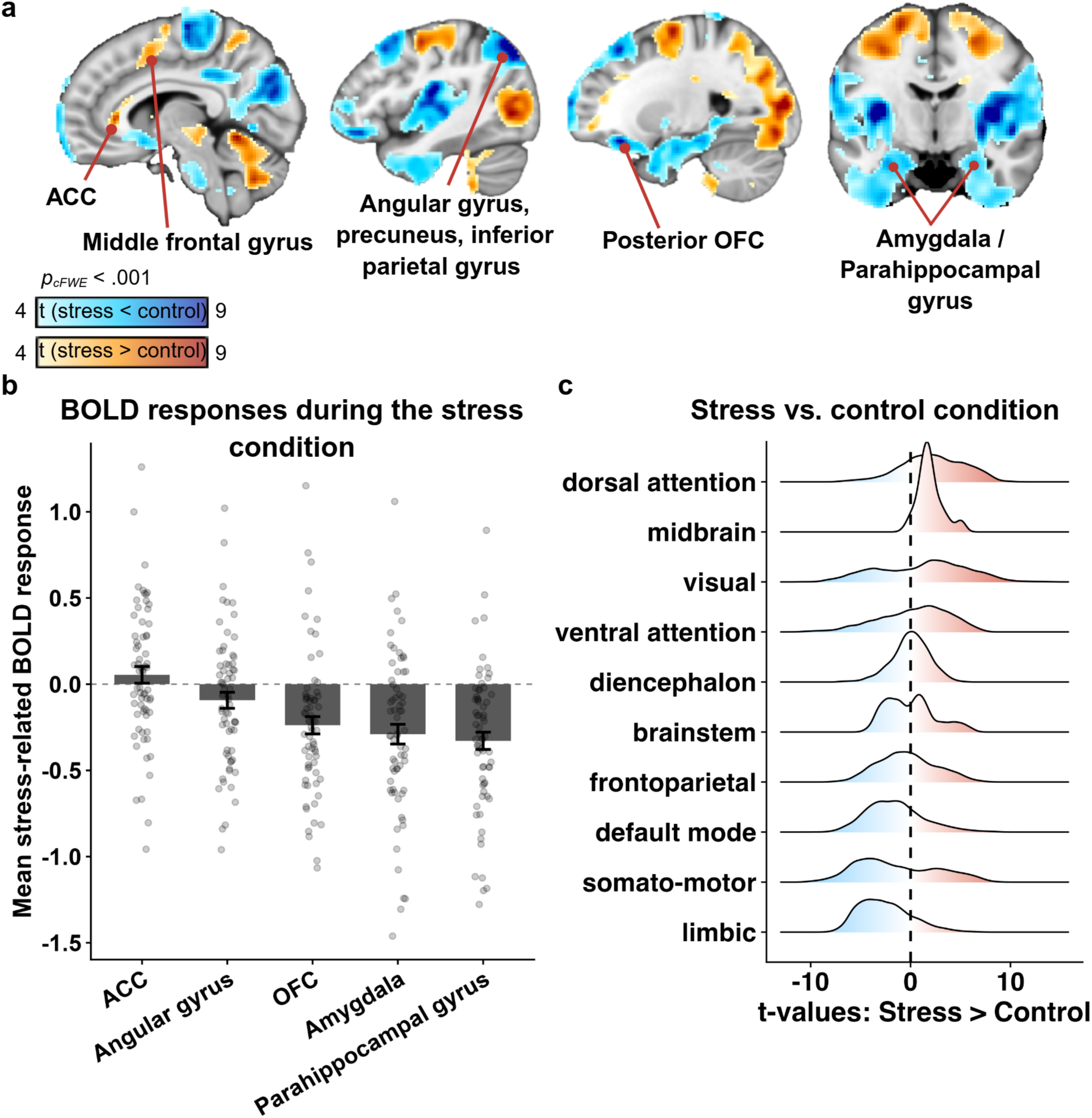
BOLD responses during the stress task. **a**: Whole-brain fMRI results show stress-related brain responses (stress vs. control). Warm colors display greater activation during stress; cool colors display higher activation during the control conditions. All regions *p* < .001 FWE cluster-corrected (FWE = family-wise error). ACC = anterior cingulate cortex. OFC = orbitofrontal cortex. **b**: For each participant, contrast estimates for stress relative to the control condition were averaged across bilateral regions defined by the Harvard-Oxford atlas. Bars represent the group means for the regions (± SEM). Individual data points show participant-level contrast estimates. The dashed horizontal line displays no difference between stress and control conditions. Negative values indicate lower BOLD responses during stress relative to the control condition. **c**: Ridgeline density plots show the distribution of grey-matter voxelwise *t*-values for the stress > control contrast. Stress-induced activation is displayed in warm colors (i.e., red) while deactivation is represented by cool colors (i.e., blue). Networks were ordered by increasing median *t*-value. The dashed vertical line indicates *t* = 0. Distributions are descriptive and suggest increased activity in the dorsal attention network and midbrain, while decreased activity is more prominent in the default mode, limbic and somato-motor networks. Figure created with BioRender.com.

### Everyday stress is associated with subjective, but not autonomic, task-related stress reactivity

Across the sample, participants reported low to moderate EMA stress levels (Figure S2a-b; *M* = 3.29 ± 1.23). To test whether everyday stress and task-induced stress reactivity are related, we predicted EMA stress with subjective task-induced stress (post minus pre stress task ratings) while controlling for mood state (satisfied minus depressed) and age, BMI and OC use. Higher task-induced stress was associated with lower EMA stress (Figure 4b; *b* = -0.13, *SE* = 0.05, *t*(63) = -2.61, *p* = .011). In contrast, the autonomic response to task-induced stress (increase in HR from baseline to the stress task) was not associated with EMA stress (*b* = -0.00, *SE* = 0.01, *t*(61) = -0.36, *p* = .720), controlling for mood state and other covariates.

**Figure 4.**
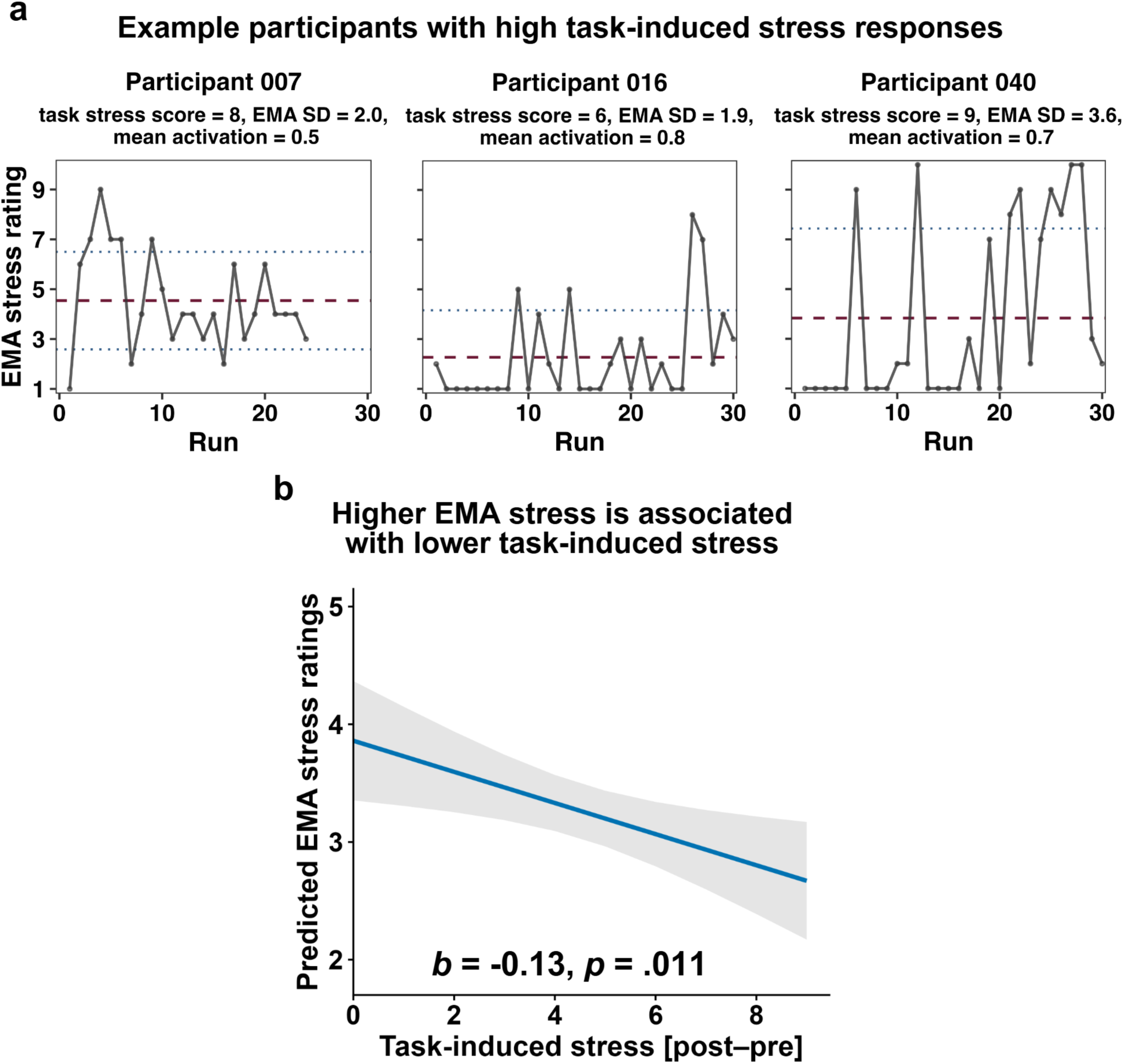
Association between EMA stress ratings and task-induced stress responses. **a:** Exemplary EMA stress trajectories illustrating participants with high task-induced stress responses (on a subjective and neural level). Task stress score = post-minus pre-stress subjective rating. Mean activation = mean stress-related brain activation within the consensus activation mask. Dashed red line indicates each participant’s mean EMA stress rating. The dotted blue lines indicate ±1 SD around the mean. **b:** Higher task stress ratings (post-minus pre-stress) were associated with lower EMA stress (*b* = -0.13, *SE* = 0.05, *t*(63) = -2.61, *p* = .011). The shaded band represents the 95% confidence interval. Ratings are adjusted for mood state, BMI, age, and OC use. Figure created with BioRender.com.

### EMA stress variability is associated with BOLD responses to acute stress

At the whole-brain level, variability in EMA stress and BOLD responses during acute stress were positively associated in the bilateral caudate (Figure 5a, S3; *t_max_* = 5.53, *k_E_* = 401, *r_ROI_* = 0.32, *p_cluster.FWE_* < .001, Table S2), cerebellum (*t_max_* = 6.02, *k_E_* = 1091, *p_cluster.FWE_* < .001), middle frontal cortex (*t_max_* = 4.76, *k_E_* = 143, *p_cluster.FWE_* = .033) and (*t_max_* = 4.74, *k_E_* = 271, *p_cluster.FWE_* < .001). In contrast, there were no significant associations with mean EMA stress (all *t_max_* < 4.01, all *p_cluster.FWE_* > .985). To examine whether the association with EMA stress variability was regionally specific or reflected a more general shift across stress-responsive regions, we next assessed mean stress-related activation within the consensus activation and deactivation masks (exemplary EMA stress trajectories of participants with high stress-related activation: Figure 4a). Mean activation within the consensus activation mask was not associated with mean EMA stress (*r* = -.07, *p* = .562, LM: *b* = -0.51, *SE* = 0.72, *t*(62) = -0.71, *p* = .483), but greater variability in EMA stress showed a robust association with higher mean stress-related activation, controlling for age, BMI and OC use (Figure 5b; *r* = .25, *p* = .041; LM: *b* = 0.73, *SE* = 0.35, *t*(62) = 2.11, *p* = .039). Mean deactivation within the consensus deactivation mask was not associated with either mean EMA stress (*r* = - .05, *p* = .714, LM: *b* = 0.21, *SE* = 0.25, *t*(62) = 0.84, *p* = .402) or EMA stress variability (*r* = .12, *p* = .340, LM: *b* = -0.21, *SE* = 0.25, *t*(62) = 0.84, *p* = .402). Consistent with the activation finding, we descriptively observed a shift towards stress-induced activation across all networks with greater EMA stress variability (Figure 5c). This association was significant after FDR correction in the dorsal attention (*b* = 0.17, *SE* = 0.06, *t*(61) = 2.69, *p_FDR_* = .031), visual (*b* = 0.25, *SE* = 0.09, *t*(61) = 2.85, *p_FDR_* = .031), frontoparietal (*b* = 0.18, *SE* = 0.07, *t*(61) = 2.47, *p_FDR_* = .041), and default mode networks (*b* = 0.19, *SE* = 0.08, *t*(61) = 2.31, *p_FDR_* = .049), as well as in the diencephalon (*b* = 0.25, *SE* = 0.09, *t*(61) = 2.69, *p_FDR_* = .031).

**Figure 5.**
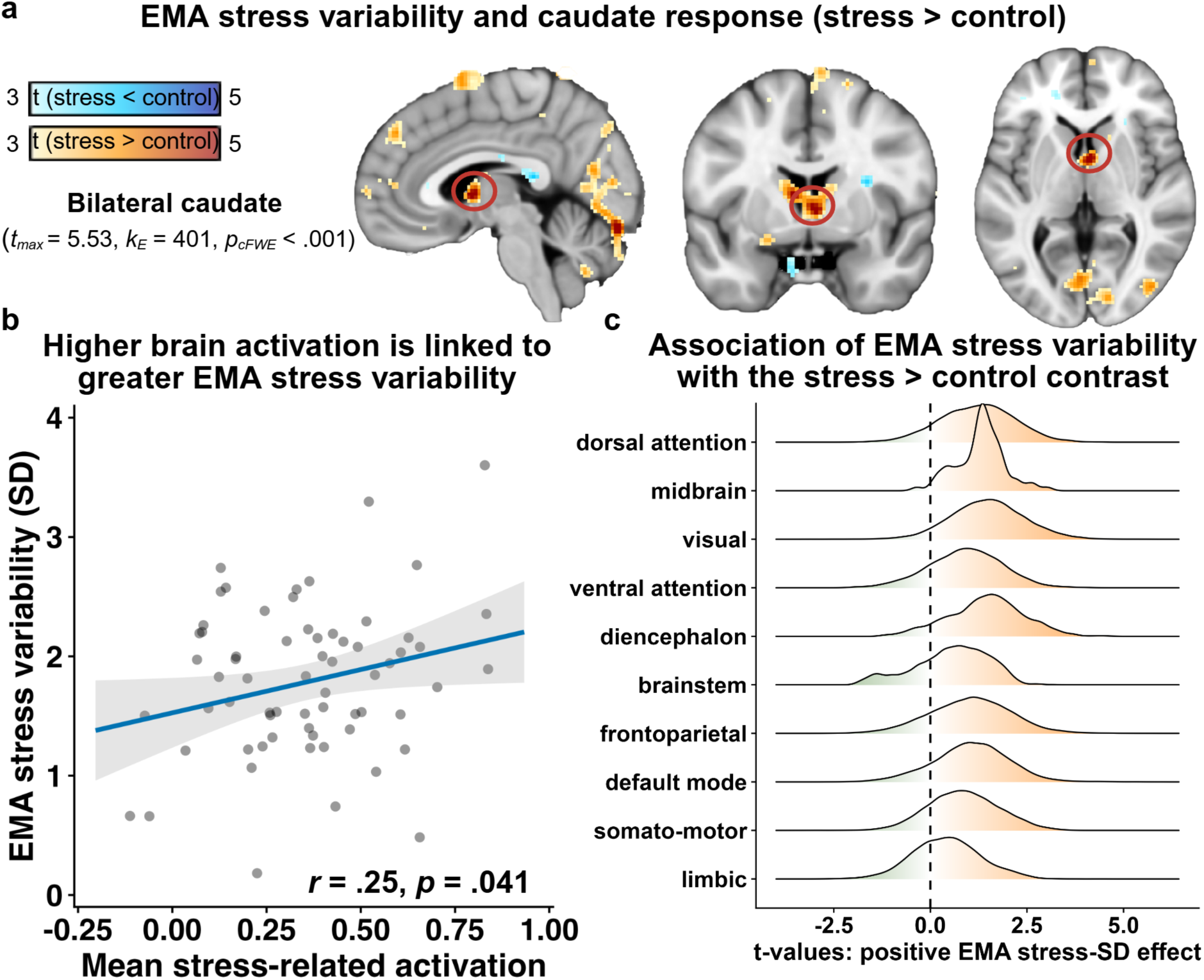
Association between variability in EMA stress and BOLD responses. **a:** Whole-brain fMRI results showed a positive association between the SD of EMA stress ratings and BOLD responses in the bilateral caudate during the stress condition (relative to the control condition; *p* < .001 FWE cluster-corrected). *k*_E_ = cluster extent. **b:** Higher mean stress-related brain activation within the consensus activation mask was associated with greater EMA stress variability (operationalized by SD; *r* = .25, *p* = .041; LM: *b* = 0.73, *p* = .039). The shaded band represents the 95% confidence interval; dots show individual values. **c**: Ridgeline density plots descriptively suggest a shift towards stress-induced activation across all networks in association with EMA stress variability. Figure created with BioRender.com.

### Association between EMA stress variability and stress-induced task-based functional connectivity

In parallel with the whole-brain BOLD-response analysis, we examined task-based functional connectivity using the bilateral vmPFC as the seed region to assess stress-induced changes in functional coupling. Compared with the control condition, stress induced stronger coupling between the vmPFC and posterior default mode regions such as the posterior cingulate cortex (*t_max_* = 18.06) and angular gyrus (*t_max_* = 17.25), as well as the dorsolateral superior frontal cortex (*t_max_* = 17.64; all *p_cluster.FWE_* < .001; Figure 6a). The cluster also extended around the vmPFC seed into the medial orbitofrontal (*t_max_* = 18.70), anterior cingulate (*t_max_* = 18.70), and medial superior frontal cortex (*t_max_* = 19.08). In contrast, coupling between the vmPFC and a cluster (Figure 6a; *k_E_* = 21581, *p_cluster.FWE_* < .001) with peaks in the right hippocampus (*t_max_* = 12.23) and the bilateral caudate was lower during stress vs. control (*t_max_* = 11.03; Table S3).

**Figure 6.**
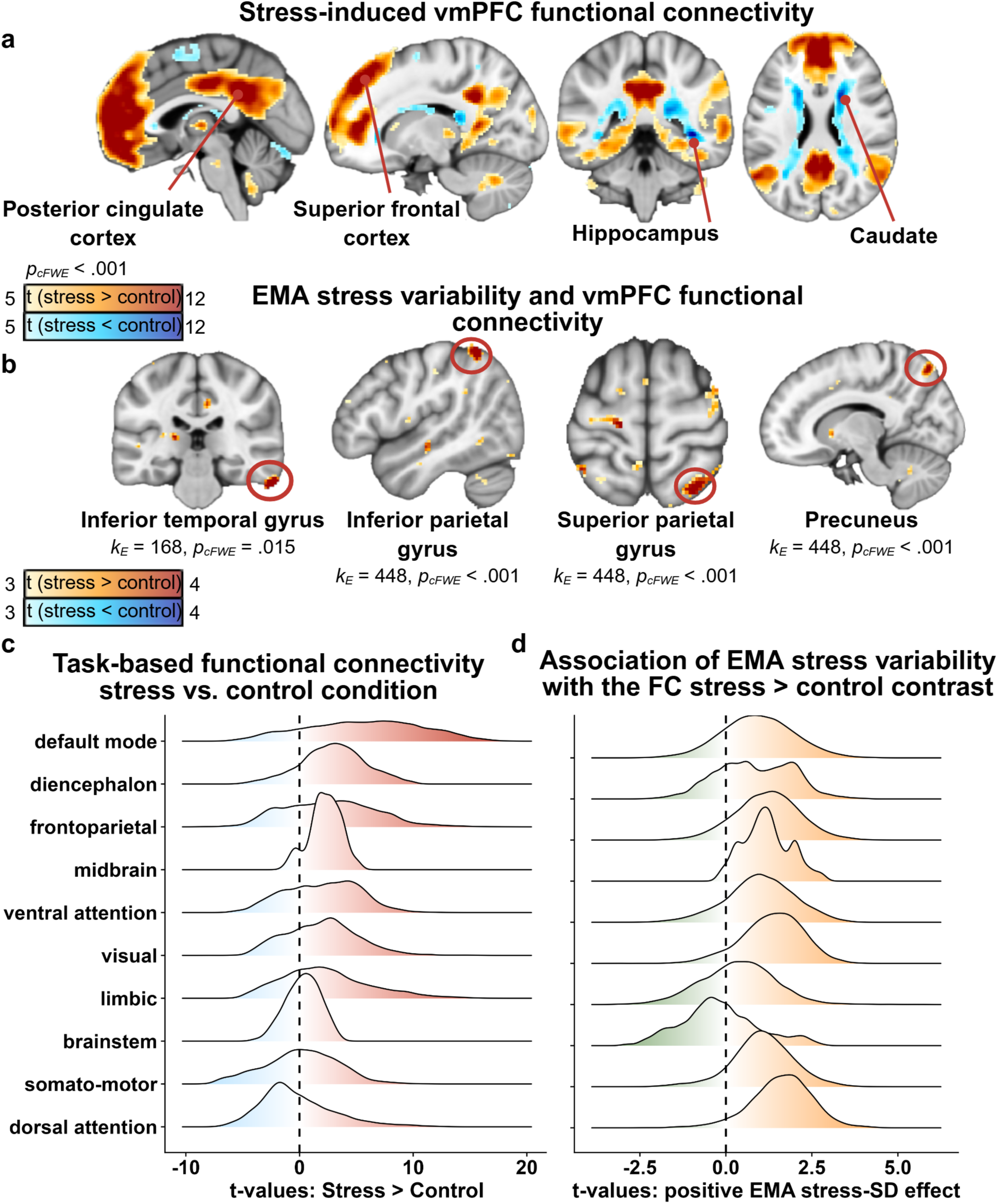
Association between EMA stress and task-based functional connectivity with the vmPFC. **a:** Task-based functional connectivity of the vmPFC during the stress and the control conditions. Warm colors (i.e., red) indicate greater vmPFC functional connectivity during stress, whereas cool colors (i.e., blue) indicate greater connectivity during the control condition. **b:** Higher functional connectivity between the vmPFC and temporal as well as parietal regions during the stress condition is positively associated with the SD of EMA stress. *p_cFWE_* = cluster-corrected family-wise error. *k_E_* = cluster extent. **c**: Ridgeline density plots descriptively suggest that vmPFC connectivity was stronger during the stress conditions (relative to the control) across most networks. **d**: Descriptively, greater variability in EMA stress was associated with a shift towards a more positive stress-related vmPFC connectivity across most networks. Figure created with BioRender.com.

We next examined whether EMA stress variability was associated not only with regional activation but also with stress-related functional connectivity (Figure 6b). During the stress condition relative to control, we observed a positive relationship between EMA stress variability and task-based vmPFC connectivity, with clusters in the inferior (*t_max_* = 4.73) and superior parietal gyrus (*t_max_* = 4.56), as well as in the precuneus (*t_max_* = 4.26, *k_E_* = 448, all *p_cluster.FWE_ <* .001) and in the right inferior temporal gyrus (*t_max_* = 3.58, *k_E_* = 149, *p_cluster.FWE_* = .025) (Table S4).

At the network level, the stress manipulation led to widespread changes in vmPFC functional connectivity. Descriptively, vmPFC connectivity was generally stronger during the stress conditions than during the control conditions across most networks. This positive shift was especially pronounced in the default mode (*t*(66) = 15.30, *p_FDR_* < .001), diencephalic (*t*(66) = 5.42, *p_FDR_* < .001), frontoparietal (*t*(66) = 5.79, *p_FDR_* < .001), limbic (*t*(66) = 6.48, *p_FDR_* < .001), midbrain (*t*(66) = 3.42, *p_FDR_* = .002), ventral attention (*t*(66) = 4.33, *p_FDR_* < .001), and visual networks (*t*(66) = 3.07, *p_FDR_* = .004). No significant changes were observed in the brainstem, dorsal attention, or somato-motor networks (all *p_FDR_* > .05; Figure 6c). Greater variability in EMA stress was associated with more positive stress-related vmPFC connectivity across most networks (Figure 6d). After FDR correction, higher EMA stress variability was significantly associated with greater stress-related vmPFC connectivity in the dorsal attention (*b* = 0.08, *SE* = 0.03, *t*(61) = 2.90, *p_FDR_* = .038) and frontoparietal networks (*b* = 0.06, *SE* = 0.02, *t*(61) = 2.76, *p_FDR_* = .038; Figure 6d) whereas associations in the default mode, somato-motor, ventral attention, visual, midbrain, diencephalic, limbic networks, and brainstem did not survive FDR correction (all *p_FDR_* > .05).

### Sensitivity analysis

#### EMA stress ratings do not differ by OC group

To test the robustness of our findings, we performed a sensitivity analysis, evaluating the effect of changes in OC use on EMA stress. We assessed whether the OC groups differed in their EMA stress levels, as they changed their OC use status after the first fMRI session. OC group did not predict mean EMA stress (*F*(2, 62) = 0.19, *p* = .831) or variability in EMA stress (*F*(2, 62) = .74, *p* = .480), controlling for confounds (Figure S2c). As OC groups did not predict any EMA stress-related measures, all presented analyses used the simpler binary OC group covariate (OC use vs. non-use) rather than the three-level OC group variable.

## Discussion

Acute laboratory stress tasks are widely used to assess individual differences in stress responses, yet it remains unclear how these acute responses relate to stress experienced in everyday life. By combining an fMRI stress paradigm with 3-month EMA in a sample of healthy young women, we observed that acute stress-induced activation and vmPFC connectivity across widespread networks were associated with variability in everyday stress but not with participants’ average stress levels. Higher overall everyday stress was also related to lower task-induced subjective stress, whereas autonomic responses to acute stress were not linked to everyday stress. Thus, acute stress responses in the brain may reflect greater reactivity to stress in daily life (i.e., larger fluctuations over time) rather than overall levels of high or low stress exposure, whereas subjective stress responses appear more closely related to overall everyday stress. Taken together, our findings suggest that different components of the acute stress response provide complementary information about everyday stress, which is important for how acute stress responses are interpreted in relation to stress exposure outside the laboratory.

In line with previous studies, we observed higher activation during stress conditions in several brain regions involved in cognitive and emotion processing, while limbic and default mode network regions showed higher activation during control conditions (Berretz et al., 2021; Kogler et al., 2015a; Noack et al., 2019). These activation changes may reflect a shift in neural processing after stress onset, redirecting attention from ongoing environmental appraisal toward the immediate stressor (Giglberger et al., 2023; Henze et al., 2020; Hermans et al., 2014; Pruessner et al., 2008; van Oort et al., 2017). We also found stress-induced changes in vmPFC functional connectivity that broadly align with previous findings of increased vmPFC connectivity with prefrontal regions and altered connectivity with posterior default mode network regions during acute stress (Ginty et al., 2019; Kühnel et al., 2022; Sinha et al., 2016). Overall, the MIST induced the expected neural effects of acute stress in our sample.

Crucially, greater variability in everyday stress across the EMA period was positively associated with a general increase in stress-induced brain activation during the MIST. Across task-positive regions, one of the more pronounced associations was observed in the caudate, which has previously been implicated in psychosocial stress (Kogler et al., 2015a; Kühnel et al., 2022). In addition, greater everyday stress variability was associated with stronger stress-induced functional connectivity between the vmPFC and posterior and parietal regions as well as with the inferior temporal gyrus. These findings fit within previous literature linking acute neural stress responses to stress processes outside the laboratory, although prior studies have focused primarily on overall daily stress or responses to stressful events. For example, acute neural stress responses have been associated with subsequent changes in perceived stress during prolonged real-life stress (Giglberger et al., 2023) and with affective reactivity to stressful events in daily life (Tutunji et al., 2025). Higher daily stress during the preceding day has also been associated with stronger connectivity between the hippocampus and the vmPFC/subgenual anterior cingulate cortex (Ren et al., 2022). Building on these findings, our results suggest that everyday stress variability may represent an additional dimension of individual differences in acute neural stress responding. This relevance is supported by studies showing that fluctuations in daily stress are linked to differences in stress-related vulnerability (Doubková et al., 2025) and by evidence that stress variability may complement mean stress levels in clinical risk assessment (Eddie et al., 2021). Greater stress fluctuations have also been related to greater emotional variability in daily life (Xia et al., 2021), and interoceptive accuracy has been linked to mood variability but not average mood (Kaduk et al., 2026). This may also be relevant from a mental health perspective, as greater affect variability has been associated with poorer concurrent and subsequent mental health outcomes (Hardy & Segerstrom, 2017; Jenkins et al., 2024). Taken together, our finding that greater everyday stress variability was associated with stronger acute neural stress responses therefore suggests that variability in everyday stress may capture individual differences in the sensitivity of the stress system to changing demands that could be relevant for vulnerability to stress-related mental health problems.

In contrast to everyday stress variability, average everyday stress was not related to acute brain responses. This may appear surprising, as acute subjective stress responses have been associated with concurrent activation changes in prefrontal and limbic regions during laboratory stress (Meier & Schwabe, 2024; Orem et al., 2019; Wheelock et al., 2016). However, these studies examine neural and subjective responses elicited by the same acute stressor, while our measure reflected average perceived stress repeatedly across a prolonged period in everyday life, limiting direct comparability. Furthermore, although autonomic responses were not associated with everyday stress, higher fasting cortisol was associated with higher HR across the stress task, consistent with previous findings linking baseline cortisol levels to cardiovascular activity during acute stress (Pico-Alfonso et al., 2007). We also observed that higher task-induced subjective stress was associated with lower everyday stress. Previous studies report mixed findings. Some show higher perceived stress over the preceding month being associated with greater acute subjective stress responses (Creswell et al., 2025), while others demonstrate associations with greater negative emotion during laboratory stress but not with a stronger acute increase (Chen et al., 2026), or no associations (Ren et al., 2022). In contrast, higher chronic or recent life stress has been associated with blunted physiological responses to acute stress (Howard et al., 2023; Xin et al., 2020), suggesting that greater ongoing stress does not necessarily correspond to stronger acute stress reactivity. Notably, these studies examined prior stress in relation to subsequent acute stress responses and are therefore not directly comparable to our design. Differences across studies may also partly reflect how stress responses are assessed and the time period covered by these measures (Weber et al., 2022). Together with the associations observed for stress-induced brain responses, these findings further suggest that different components of the acute stress response provide complementary information about stress experienced outside the laboratory.

While our study is one of the few to combine acute laboratory stress with EMA, some limitations should be considered. First, we did not assess whether participants encountered unusual or specific stressors during the EMA period. We therefore cannot determine whether the observed associations differ depending on stressor type, intensity, or context. Second, coping strategies were not accounted for, although they may influence responses to and regulation of everyday stressors (O’Rourke et al., 2022; Sun et al., 2023). Third, despite the extensive EMA period, perceived stress was assessed only once in the evening on prompted days, potentially limiting our ability to capture within-day fluctuations. Future studies could address this by using wearable devices that continuously monitor physiological signals, such as HR or cortisol levels (Ding et al., 2024; Tu et al., 2025), to trigger EMA assessments when increases in these signals are detected. This could allow stress responses to be captured closer to their occurrence. Finally, generalizability is limited by the inclusion of healthy young women only, as the study was initially conducted to investigate OC use. Given reported sex differences in neural stress responses (Cohen et al., 2023; Kogler et al., 2015b; Kühnel et al., 2023), the findings may not generalize to men or older populations. However, female-only samples have been underrepresented in acute social stress research (Rrapaj et al., 2023) and women’s reproductive health factors, including OC use, closing an important gap in human neuroimaging (Taylor et al., 2021). Characterizing stress-related processes in women is also clinically relevant given the higher prevalence and burden of several stress-related disorders, including PTSD and depression, in women than in men (Haering et al., 2024; Salk et al., 2017).

Understanding how experimentally induced stress relates to stress experienced in everyday life is important for determining what laboratory stress paradigms capture about individual differences in stress responding. Here, combining an fMRI stress paradigm with EMA, we demonstrate that acute stress-related brain responses were associated with variability, but not average levels, of perceived everyday stress. Crucially, these associations extended beyond regional brain responses to stress-induced changes in vmPFC connectivity. Our findings therefore highlight everyday stress variability as an important aspect when linking experimentally induced stress to naturalistic stress experiences. A better understanding of which aspects of everyday stress experience are reflected in laboratory stress responses may ultimately help clarify the relevance of acute stress paradigms for investigating individual differences in stress-related vulnerability.

## Acknowledgement

We thank Ulrike Bernig, Judith Scharpf, Lisa Stark and Grete Moeller for their help with data acquisition. This study was conducted as part of the International Research Training Group: Women’s Mental Health Across the Reproductive Years (IRTG 2804). MK, SB and ASK were supported by IRTG 2804, funded by the Deutsche Forschungsgemeinschaft (DFG, German Research Foundation); grant number: GRK 2804/1. In addition, NBK was supported by DFG grants KR 4555/7-1; KR 4555/9-1; and KR 4555/10-1. AK received support by the Else Kröner-Fresenius Stiftung, grant 2024_EKEA.149.

## Author contributions

NBK, MH, ISP and MK were responsible for the study concept and design. ASK and LK contributed to study planning and implementation. MK collected data under supervision by NBK and MH. NBK conceived the method of this paper and AK and MK processed the data. MK performed the data analysis and NBK and AK contributed to analyses. MK, AK and NBK wrote the manuscript. All authors contributed to the interpretation of findings, provided critical revision of the manuscript for important intellectual content and approved the final version for publication.

## Financial disclosure

The authors declare no competing financial interests.

## Notes

https://github.com/neuromadlab/Stress_EMA_fMRI

### Competing Interest Statement

The authors have declared no competing interest.

## References

Albert, K., Pruessner, J., & Newhouse, P. (2015). Estradiol levels modulate brain activity and negative responses to psychosocial stress across the menstrual cycle. Psychoneuroendocrinology, 59, 14–24. 10.1016/j.psyneuen.2015.04.022

Barel, E., Abu-Shkara, R., Colodner, R., Masalha, R., Mahagna, L., Zemel, O. C., & Cohen, A. (2018). Gonadal hormones modulate the HPA-axis and the SNS in response to psychosocial stress. Journal of Neuroscience Research, 96(8), 1388–1397. 10.1002/jnr.24259

Berretz, G., Packheiser, J., Kumsta, R., Wolf, O. T., & Ocklenburg, S. (2021). The brain under stress—A systematic review and activation likelihood estimation meta-analysis of changes in BOLD signal associated with acute stress exposure. Neuroscience & Biobehavioral Reviews, 124, 89–99. 10.1016/j.neubiorev.2021.01.001

BIOPAC Systems, Inc. (2024). *AcqKnowledge software* (Version 5.0.8) [Computer software]. https://www.biopac.com/product/acqknowledge-software/

Bürger, Z., Kordowich, C., Kübbeler, J., Müllerschön, C., Kimmig, A.-C. S., Su, M., Lämmerhofer, M., Sacher, J., Henes, M., Comasco, E., Derntl, B., & Kogler, L. (2025). Subjective, behavioural and physiological correlates of stress in women using hormonal contraceptives. The British Journal of Psychiatry, 226(6), 392–400. 10.1192/bjp.2025.7

Chen, M., Gao, M., Shao, R., Tong, H., Liu, J. M., Cheung, A. K., & Lee, T. M. C. (2026). Chronic stress modulates the relationship between acute stress-related cortical-limbic circuit functional connectivity and depression symptoms. Journal of Affective Disorders, 395, 120725. 10.1016/j.jad.2025.120725

Chrousos, G. P. (1992). The Concepts of Stress and Stress System Disorders: Overview of Physical and Behavioral Homeostasis. JAMA, 267(9), 1244. 10.1001/jama.1992.03480090092034

Constantin, A.-E., & Patil, I. (2021). ggsignif: R Package for Displaying Significance Brackets for ‘ggplot2’. PsyArxiv. 10.31234/osf.io/7awm6

Creswell, D., Brown, K. W., Cohen, S., Creswell, K., Zoccola, P., Dickerson, S., Dutcher, J., Wu, S., & Chin, B. (2025). Does high perceived stress over the past month alter cortisol reactivity to the Trier Social Stress Test? Psychoneuroendocrinology, 172, 107256. 10.1016/j.psyneuen.2024.107256

De Calheiros Velozo, J., Vaessen, T., Lafit, G., Claes, S., & Myin-Germeys, I. (2023). Is daily-life stress reactivity a measure of stress recovery? An investigation of laboratory and daily-life stress. Stress and Health, 39(3), 638–650. 10.1002/smi.3213

Dedovic, K., Renwick, R., Mahani, N. K., Engert, V., Lupien, S. J., & Pruessner, J. C. (2005). The Montreal Imaging Stress Task: Using functional imaging to investigate the effects of perceiving and processing psychosocial stress in the human brain. Journal of Psychiatry and Neuroscience, 30(5), 319–325.

Desikan, R. S., Ségonne, F., Fischl, B., Quinn, B. T., Dickerson, B. C., Blacker, D., Buckner, R. L., Dale, A. M., Maguire, R. P., Hyman, B. T., Albert, M. S., & Killiany, R. J. (2006). An automated labeling system for subdividing the human cerebral cortex on MRI scans into gyral based regions of interest. NeuroImage, 31(3), 968–980. 10.1016/j.neuroimage.2006.01.021

Ding, Y., Tan, K., Sheng, L., Ren, H., Su, Z., Yang, H., Zhang, X., Li, J., & Hu, P. (2024). Integrated mental stress smartwatch based on sweat cortisol and HRV sensors. Biosensors and Bioelectronics, 265, 116691. 10.1016/j.bios.2024.116691

Doubková, N., Zlámal, F., Fňašková, M., Preiss, M., Nečasová, M., Wolframová, N., Svoboda, V., Ulčák, D., & Rektor, I. (2025). Daily Stress Variability in Two Generations of Survivors of the War in the Former Yugoslavia. Stress and Health, 41(5), e70113. 10.1002/smi.70113

Eddie, D., Barr, M., Njeim, L., & Emery, N. (2021). Mean Versus Variability: Disentangling Stress Effects on Alcohol Lapses Among Individuals in the First Year of Alcohol Use Disorder Recovery. Journal of Studies on Alcohol and Drugs, 82(5), 623–628. 10.15288/jsad.2021.82.623

Elbau, I. G., Brücklmeier, B., Uhr, M., Arloth, J., Czamara, D., Spoormaker, V. I., Czisch, M., Stephan, K. E., Binder, E. B., & Sämann, P. G. (2018). The brain’s hemodynamic response function rapidly changes under acute psychosocial stress in association with genetic and endocrine stress response markers. Proceedings of the National Academy of Sciences, 115(43), E10206–E10215. 10.1073/pnas.1804340115

Esteban, O., Markiewicz, C. J., Blair, R. W., Moodie, C. A., Isik, A. I., Erramuzpe, A., Kent, J. D., Goncalves, M., DuPre, E., Snyder, M., Oya, H., Ghosh, S. S., Wright, J., Durnez, J., Poldrack, R. A., & Gorgolewski, K. J. (2019). fMRIPrep: A robust preprocessing pipeline for functional MRI. Nature Methods, 16(1), 111–116. 10.1038/s41592-018-0235-4

Giglberger, M., Peter, H. L., Henze, G.-I., Kraus, E., Bärtl, C., Konzok, J., Kreuzpointner, L., Kirsch, P., Kudielka, B. M., & Wüst, S. (2023). Neural responses to acute stress predict chronic stress perception in daily life over 13 months. Scientific Reports, 13(1), 19990. 10.1038/s41598-023-46631-w

Ginty, A. T., Kraynak, T. E., Kuan, D. C., & Gianaros, P. J. (2019). Ventromedial prefrontal cortex connectivity during and after psychological stress in women. Psychophysiology, 56(11), e13445. 10.1111/psyp.13445

Habib, K. E., Gold, P. W., & Chrousos, G. P. (2001). NEUROENDOCRINOLOGY OF STRESS. Endocrinology and Metabolism Clinics of North America, 30(3), 695–728. 10.1016/S0889-8529(05)70208-5

Haering, S., Schulze, L., Geiling, A., Meyer, C., Klusmann, H., Schumacher, S., Knaevelsrud, C., & Engel, S. (2024). Higher risk—less data: A systematic review and meta-analysis on the role of sex and gender in trauma research. Journal of Psychopathology and Clinical Science, 133(3), 257–272. 10.1037/abn0000899

Hardy, J., & Segerstrom, S. C. (2017). Intra-individual variability and psychological flexibility: Affect and health in a National US sample. Journal of Research in Personality, Within-Person Variability in Personality, 69, 13–21. 10.1016/j.jrp.2016.04.002

Henze, G.-I., Konzok, J., Kreuzpointner, L., Bärtl, C., Peter, H., Giglberger, M., Streit, F., Kudielka, B. M., Kirsch, P., & Wüst, S. (2020). Increasing Deactivation of Limbic Structures Over Psychosocial Stress Exposure Time. Biological Psychiatry: Cognitive Neuroscience and Neuroimaging, 5(7), 697–704. 10.1016/j.bpsc.2020.04.002

Herman, J. P., Ostrander, M. M., Mueller, N. K., & Figueiredo, H. (2005). Limbic system mechanisms of stress regulation: Hypothalamo-pituitary-adrenocortical axis. *Progress in Neuro-Psychopharmacology and Biological Psychiatry*, Experimental Stress: From Basic to Clinical Aspects, 29(8), 1201–1213. 10.1016/j.pnpbp.2005.08.006

Hermans, E. J., Henckens, M. J. A. G., Joëls, M., & Fernández, G. (2014). Dynamic adaptation of large-scale brain networks in response to acute stressors. Trends in Neurosciences, 37(6), 304–314. 10.1016/j.tins.2014.03.006

Hossein, S., Cooper, J. A., DeVries, B. A. M., Nuutinen, M. R., Hahn, E. C., Kragel, P. A., & Treadway, M. T. (2023). Effects of acute stress and depression on functional connectivity between prefrontal cortex and the amygdala. Molecular Psychiatry, 28(11), 4602–4612. 10.1038/s41380-023-02056-5

Howard, S., Gallagher, S., Ginty, A. T., & Whittaker, A. C. (2023). Life event stress is associated with blunted cardiovascular responding to both personally salient and personally non-salient laboratory tasks. Psychophysiology, 60(3), e14199. 10.1111/psyp.14199

Jenkins, B. N., Ong, L. Q., Ong, A. D., Lee, H. Y. (Helen), & Boehm, J. K. (2024). Mean Affect Moderates the Association between Affect Variability and Mental Health. Affective Science, 5(2), 99–114. 10.1007/s42761-024-00238-0

Kaduk, K., Kaeber, M., Kühnel, A., Torrado, M. B., Grahlow, M., Derntl, B., & Kroemer, N. B. (2026). Glucose levels are associated with mood, but the association is mediated by ratings of metabolic state. eBioMedicine, 124. 10.1016/j.ebiom.2025.106035

Kay, M. (2024). ggdist: Visualizations of Distributions and Uncertainty in the Grammar of Graphics. IEEE Transactions on Visualization and Computer Graphics, 30(1), 414– 424. 10.1109/TVCG.2023.3327195

Kay, M. (2025). ggdist: Visualizations of Distributions and Uncertainty. 10.5281/zenodo.3879620

Kogler, L., Müller, V. I., Chang, A., Eickhoff, S. B., Fox, P. T., Gur, R. C., & Derntl, B. (2015). Psychosocial versus physiological stress—Meta-analyses on deactivations and activations of the neural correlates of stress reactions. NeuroImage, 119, 235–251. 10.1016/j.neuroimage.2015.06.059

Kördel, M., Meier, M., Kühnel, A., & Kroemer, N. B. (2025). Metabolic state shapes cortisol reactivity to acute stress: A systematic review and meta-analysis of metabolic and hormonal modulators. Neurobiology of Stress, 39, 100764. 10.1016/j.ynstr.2025.100764

Kubios Oy. (2023). *Kubios HRV* (Version 4.1.2.1) [Computer software]. https://www.kubios.com/

Kuhn, L., Noack, H., Wagels, L., Prothmann, A., Schulik, A., Aydin, E., Nieratschker, V., Derntl, B., & Habel, U. (2023). Sex-dependent multimodal response profiles to psychosocial stress. Cerebral Cortex, 33(3), 583–596. 10.1093/cercor/bhac086

Kühnel, A., Czisch, M., Sämann, P. G., Brückl, T., Spoormaker, V. I., Erhardt, A., Grandi, N. C., Ziebula, J., Elbau, I. G., Namendorf, T., Lucae, S., Binder, E. B., & Kroemer, N. B. (2022). Spatiotemporal Dynamics of Stress-Induced Network Reconfigurations Reflect Negative Affectivity. Biological Psychiatry, Molecular Mechanisms of Resilience and Vulnerability to Stress, 92(2), 158–169. 10.1016/j.biopsych.2022.01.008

Kühnel, A., Kroemer, N. B., Elbau, I. G., Czisch, M., Sämann, P. G., Walter, M., Group, B. working, & Binder, E. B. (2020). Psychosocial stress reactivity habituates following acute physiological stress. Human Brain Mapping, 41(14), 4010–4023. 10.1002/hbm.25106

Kuznetsova, A., Brockhoff, P. B., & Christensen, R. H. B. (2017). lmerTest Package: Tests in Linear Mixed Effects Models. Journal of Statistical Software, 82(13), 1–26. 10.18637/jss.v082.i13

Lenth, R. V., & Piaskowski, J. (2025). emmeans: Estimated Marginal Means, aka Least-Squares Means. 10.32614/CRAN.package.emmeans

McLaren, D. G., Ries, M. L., Xu, G., & Johnson, S. C. (2012). A generalized form of context-dependent psychophysiological interactions (gPPI): A comparison to standard approaches. NeuroImage, 61(4), 1277–1286. 10.1016/j.neuroimage.2012.03.068

Meier, J. K., & Schwabe, L. (2024). Consistently increased dorsolateral prefrontal cortex activity during the exposure to acute stressors. Cerebral Cortex, 34(4), bhae159. 10.1093/cercor/bhae159

Ming, Q., Zhong, X., Zhang, X., Pu, W., Dong, D., Jiang, Y., Gao, Y., Wang, X., Detre, J. A., Yao, S., & Rao, H. (2017). State-Independent and Dependent Neural Responses to Psychosocial Stress in Current and Remitted Depression. American Journal of Psychiatry, 174, 971–979. 10.1176/appi.ajp.2017.16080974

Müller, S. J., Teckentrup, V., Rebollo, I., Hallschmid, M., & Kroemer, N. B. (2022). Vagus nerve stimulation increases stomach-brain coupling via a vagal afferent pathway. Brain Stimulation, 15(5), 1279–1289. 10.1016/j.brs.2022.08.019

Noack, H., Nolte, L., Nieratschker, V., Habel, U., & Derntl, B. (2019). Imaging stress: An overview of stress induction methods in the MR scanner. Journal of Neural Transmission, 126(9), 1187–1202. 10.1007/s00702-018-01965-y

Orem, T. R., Wheelock, M. D., Goodman, A. M., Harnett, N. G., Wood, K. H., Gossett, E. W., Granger, D. A., Mrug, S., & Knight, D. C. (2019). Amygdala and prefrontal cortex activity varies with individual differences in the emotional response to psychosocial stress. Behavioral Neuroscience, 133(2), 203–211. 10.1037/bne0000305

O’Rourke, T., Vogel, C., John, D., Pryss, R., Schobel, J., Haug, F., Haug, J., Pieh, C., Nater, U. M., Feneberg, A. C., Reichert, M., & Probst, T. (2022). The Impact of Coping Styles and Gender on Situational Coping: An Ecological Momentary Assessment Study With the mHealth Application TrackYourStress. Frontiers in Psychology, 13. 10.3389/fpsyg.2022.913125

Pauli, W. M., Nili, A. N., & Tyszka, J. M. (2018). A high-resolution probabilistic in vivo atlas of human subcortical brain nuclei. Scientific Data, 5(1), 180063. 10.1038/sdata.2018.63

Pico-Alfonso, M. A., Mastorci, F., Ceresini, G., Ceda, G. P., Manghi, M., Pino, O., Troisi, A., & Sgoifo, A. (2007). Acute psychosocial challenge and cardiac autonomic response in women: The role of estrogens, corticosteroids, and behavioral coping styles. Psychoneuroendocrinology, 32(5), 451–463. 10.1016/j.psyneuen.2007.02.009

Posit team. (2026). RStudio: Integrated Development Environment for R. Posit Software, PBC. http://www.posit.co/

Pruessner, J. C., Dedovic, K., Khalili-Mahani, N., Engert, V., Pruessner, M., Buss, C., Renwick, R., Dagher, A., Meaney, M. J., & Lupien, S. (2008). Deactivation of the Limbic System During Acute Psychosocial Stress: Evidence from Positron Emission Tomography and Functional Magnetic Resonance Imaging Studies. Biological Psychiatry, 63(2), 234– 240. 10.1016/j.biopsych.2007.04.041

Ren, X., Zhao, X., Li, J., Liu, Y., Ren, Y., Pruessner, J. C., & Yang, J. (2022). The Hippocampal–Ventral Medial Prefrontal Cortex Neurocircuitry Involvement in the Association of Daily Life Stress With Acute Perceived Stress and Cortisol Responses. Psychosomatic Medicine, 84(3), 276–287. 10.1097/PSY.0000000000001058

Research Imaging Institute, UTHSCSA. (n.d.). *Mango: Multi-image Analysis GUI* (Version 4.1) [Computer software]. Research Imaging Institute, University of Texas Health Science Center at San Antonio. Retrieved https://mangoviewer.com/

Roy, M., Shohamy, D., & Wager, T. D. (2012). Ventromedial prefrontal-subcortical systems and the generation of affective meaning. Trends in Cognitive Sciences, 16(3), 147– 156. 10.1016/j.tics.2012.01.005

Rrapaj, A., Landau, A. M., & Winterdahl, M. (2023). Exploration of possible sex bias in acute social stress research: A semi-systematic review. Acta Neuropsychiatrica, 35(4), 205– 217. 10.1017/neu.2023.16

Salk, R. H., Hyde, J. S., & Abramson, L. Y. (2017). Gender differences in depression in representative national samples: Meta-analyses of diagnoses and symptoms. Psychological Bulletin, 143(8), 783–822. 10.1037/bul0000102

Sapolsky, R. M., Romero, L. M., & Munck, A. U. (2000). How Do Glucocorticoids Influence Stress Responses? Integrating Permissive, Suppressive, Stimulatory, and Preparative Actions*. Endocrine Reviews, 21(1), 55–89. 10.1210/edrv.21.1.0389

Sharma, R., Smith, S. A., Boukina, N., Dordari, A., Mistry, A., Taylor, B. C., Felix, N., Cameron, A., Fang, Z., Smith, A., & Ismail, N. (2020). Use of the birth control pill affects stress reactivity and brain structure and function. Hormones and Behavior, 124, 104783. 10.1016/j.yhbeh.2020.104783

Sinha, R., Lacadie, C. M., Constable, R. T., & Seo, D. (2016). Dynamic neural activity during stress signals resilient coping. Proceedings of the National Academy of Sciences, 113(31), 8837–8842. 10.1073/pnas.1600965113

Sun, T., Yap, Y., Tung, Y. C., Bei, B., & Wiley, J. F. (2023). Coping strategies predict daily emotional reactivity to stress: An ecological momentary assessment study. Journal of Affective Disorders, 332, 309–317. 10.1016/j.jad.2023.03.090

Taylor, C. M., Pritschet, L., & Jacobs, E. G. (2021). The scientific body of knowledge – Whose body does it serve? A spotlight on oral contraceptives and women’s health factors in neuroimaging. Frontiers in Neuroendocrinology, 60, 100874. 10.1016/j.yfrne.2020.100874

Teckentrup, V., Krylova, M., Jamalabadi, H., Neubert, S., Neuser, M. P., Hartig, R., Fallgatter, A. J., Walter, M., & Kroemer, N. B. (2021). Brain signaling dynamics after vagus nerve stimulation. NeuroImage, 245, 118679. 10.1016/j.neuroimage.2021.118679

Tu, J., Yeom, J., Ulloa, J. C., Solomon, S. A., Min, J., Heng, W., Kim, G., Dao, J., Vemu, R., Pang, M., Wang, C., Kim, D.-H., & Gao, W. (2025). Stressomic: A wearable microfluidic biosensor for dynamic profiling of multiple stress hormones in sweat. Science Advances, 11(32), eadx6491. 10.1126/sciadv.adx6491

Tutunji, R., Krentz, M., Kogias, N., Voogd, L. de, Krause, F., Vassena, E., & Hermans, E. J. (2025). Changes in large-scale neural networks under stress are linked to affective reactivity to stress in real life. eLife, 14. 10.7554/eLife.102574.1

Vaisvaser, S., Modai, S., Farberov, L., Lin, T., Sharon, H., Gilam, A., Volk, N., Admon, R., Edry, L., Fruchter, E., Wald, I., Bar-Haim, Y., Tarrasch, R., Chen, A., Shomron, N., & Hendler, T. (2016). Neuro-Epigenetic Indications of Acute Stress Response in Humans: The Case of MicroRNA-29c. PLOS ONE, 11(1), e0146236. 10.1371/journal.pone.0146236

van Oort, J., Tendolkar, I., Hermans, E. J., Mulders, P. C., Beckmann, C. F., Schene, A. H., Fernández, G., & van Eijndhoven, P. F. (2017). How the brain connects in response to acute stress: A review at the human brain systems level. Neuroscience & Biobehavioral Reviews, 83, 281–297. 10.1016/j.neubiorev.2017.10.015

Weber, J., Angerer, P., & Apolinário-Hagen, J. (2022). Physiological reactions to acute stressors and subjective stress during daily life: A systematic review on ecological momentary assessment (EMA) studies. PLOS ONE, 17(7), e0271996. 10.1371/journal.pone.0271996

Wheelock, M. D., Harnett, N. G., Wood, K. H., Orem, T. R., Granger, D. A., Mrug, S., & Knight, D. C. (2016). Prefrontal Cortex Activity Is Associated with Biobehavioral Components of the Stress Response. Frontiers in Human Neuroscience, 10. 10.3389/fnhum.2016.00583

Wickham, H. (2016). ggplot2: Elegant Graphics for Data Analysis. Springer-Verlag New York. https://ggplot2.tidyverse.org

Xia, W., Li, L. M. W., Jiang, D., & Liu, S. (2021). Dynamics of stress and emotional experiences during COVID-19: Results from two 14-day daily diary studies. International Journal of Stress Management, 28(4), 256–265. 10.1037/str0000234

Xin, Y., Yao, Z., Wang, W., Luo, Y., Aleman, A., & Wu, J. (2020). Recent life stress predicts blunted acute stress response and the role of executive control. Stress, 23(3), 359– 367. 10.1080/10253890.2019.1687684

Yeo, B. T. T., Krienen, F. M., Sepulcre, J., Sabuncu, M. R., Lashkari, D., Hollinshead, M., Roffman, J. L., Smoller, J. W., Zöllei, L., Polimeni, J. R., Fischl, B., Liu, H., & Buckner, R. L. (2011). The organization of the human cerebral cortex estimated by intrinsic functional connectivity. Journal of Neurophysiology, 106(3), 1125–1165. 10.1152/jn.00338.2011

